# An autonomous LLM-agent platform for computational binder design and conjugation-aware prioritization of antibody–drug conjugates

**DOI:** 10.64898/2026.04.21.719907

**Authors:** Ganggang Liu, Mingjie He, Liang Sun, Fuxian Chen, Yu Zhang

**Affiliations:** Suzhou Kai Zhi Yuan Technology Co., Ltd., 218 Sangtian Street, Suzhou Industrial Park, Suzhou 215123, China; School of Life Sciences, Suzhou Medical College, Soochow University, 199 Ren’ai Road, Suzhou 215123, China; Jiangxi Provincial People’s Hospital, The First Affiliated Hospital of Nanchang Medical College, Nanchang 330006, China; Department of General Surgery, The First Affiliated Hospital of Soochow University, Suzhou 215006, China

## Abstract

Large language model (LLM) agents have automated tool use in chemistry, but orchestrating multi-step computational biology workflows—spanning structure prediction, protein design, and covalent conjugation—remains manually intensive. Here we present Open Intelligence Hub (OIH), an autonomous LLM-agent platform that dynamically plans and executes 32 containerised tools for protein binder design and antibody–drug conjugate (ADC) prioritization. OIH introduces tier-based decision routing, ipSAE-guided interface filtering, and failure-to-knowledge distillation from 265 curated cases. Across five oncology targets, the agent correctly classified all five evaluated targets and required human correction for hotspot selection in only one case, producing binders ranked by ipSAE (Nectin-4 ipTM = 0.87, HER2 ipTM = 0.85). A controlled ablation suggests that the agent’s PPI-informed routing yields improved downstream ipTM and ipSAE scores than epitope-guided alternatives. The LLM-agnostic architecture enables deployment with local or commercial models without pipeline changes. All results are computational predictions awaiting experimental validation.

## Introduction

LLM-based agents that select and invoke external tools have emerged as a paradigm for automating multi-step scientific workflows. In chemistry, ChemCrow^12^ demonstrated that GPT-4 with 18 tools could plan small-molecule synthesis; LIDDiA^13^ and DrugAgent^14^ extended this to ligand-based drug discovery. These systems share a common architecture—an LLM reads tool descriptions, plans execution order, parses intermediate outputs, and iterates until a goal is reached—but build on foundational work in LLM tool use (including ReAct reasoning, Toolformer, and function-calling frameworks) and have been largely limited to small-molecule chemistry, typically involving 5–20 lightweight tools.

Computational protein biology presents fundamentally different orchestration challenges. A typical binder design campaign requires structure retrieval, surface analysis, backbone generation (RFdiffusion^6^), sequence design (ProteinMPNN^7^), structural validation (AlphaFold 3^23^), and downstream conjugation chemistry—each running in isolated GPU containers with distinct input/output formats and multi-gigabyte memory requirements. The decision space is substantially more complex and higher-dimensional: the agent must classify targets by available structural information, select appropriate hotspot prediction methods, manage GPU memory across tools requiring 4–40 GB VRAM, and apply quality-control metrics that differ from those used in small-molecule docking. Recent binder design tools including BindCraft^8^ (10–100% experimental hit rates), AlphaProteo^9^, and PXDesign^10^ achieve impressive success rates but require expert-specified inputs—target preparation, hotspot selection, and validation thresholds—that are themselves multi-step decision problems.

Here we present OIH, an autonomous agent platform that accepts a target protein name in natural language and dynamically orchestrates 32 containerised tools across 15 Docker containers to execute the full binder design and conjugation-aware prioritization pipeline. Unlike hardcoded workflow managers (for example, ProteinDJ^19^ on Nextflow), OIH plans execution order at runtime from tool descriptions and domain knowledge injected as contextual skills. We evaluate the agent on five oncology targets spanning two decision tiers, quantify its unaided decision accuracy, and demonstrate a failure-to-knowledge distillation scheme that converts 265 execution failures into reusable agent context. The orchestration layer is LLM-agnostic: switching from local Qwen3-14B to a commercial API requires changing only the inference endpoint. We make three methodological contributions: (1) a tier-based decision routing framework that selects hotspot prediction methods based on structural priors; (2) a failure-to-knowledge distillation mechanism that converts execution failures into reusable agent context without model retraining; and (3) an empirical evaluation quantifying how agent-level upstream decisions propagate to downstream design quality.

A key methodological question we address is whether an LLM agent can make consequential upstream decisions—specifically, hotspot selection for binder design—that materially influence downstream computational outcomes. Through a controlled ablation on CD36 using three different hotspot sources under otherwise identical downstream tools, we provide a direct quantitative comparison showing that the agent’s routing decision is not merely a convenience but a critical variable. We further introduce ipSAE, a recently proposed interface metric, as an automated quality gate that detects designs lacking genuine interface contacts—a failure mode invisible to the standard ipTM score.

## Results

### Agent architecture and dynamic planning

OIH comprises three layers (Fig. 1a): a natural-language interface, a Qwen3-14B orchestration engine (4-bit AWQ quantisation, vLLM^41^ serving), and a containerised tool ecosystem. The 32 tools span six categories: structure prediction (AlphaFold 3^23^, ESM2^31^, IgFold^32^, PeSTo^16^), protein design (RFdiffusion^6^, ProteinMPNN^7^, BindCraft^8^), molecular docking (GNINA^33^, AutoDock-GPU^34^, DiffDock^35^), analysis (fpocket^36^, P2Rank^37^, FreeSASA^38^, GROMACS^39^), immunology (DiscoTope3^15^), and chemistry (Chemprop^40^, RDKit). Full tool versions are listed in Supplementary Table S1.

**Fig. 1.**
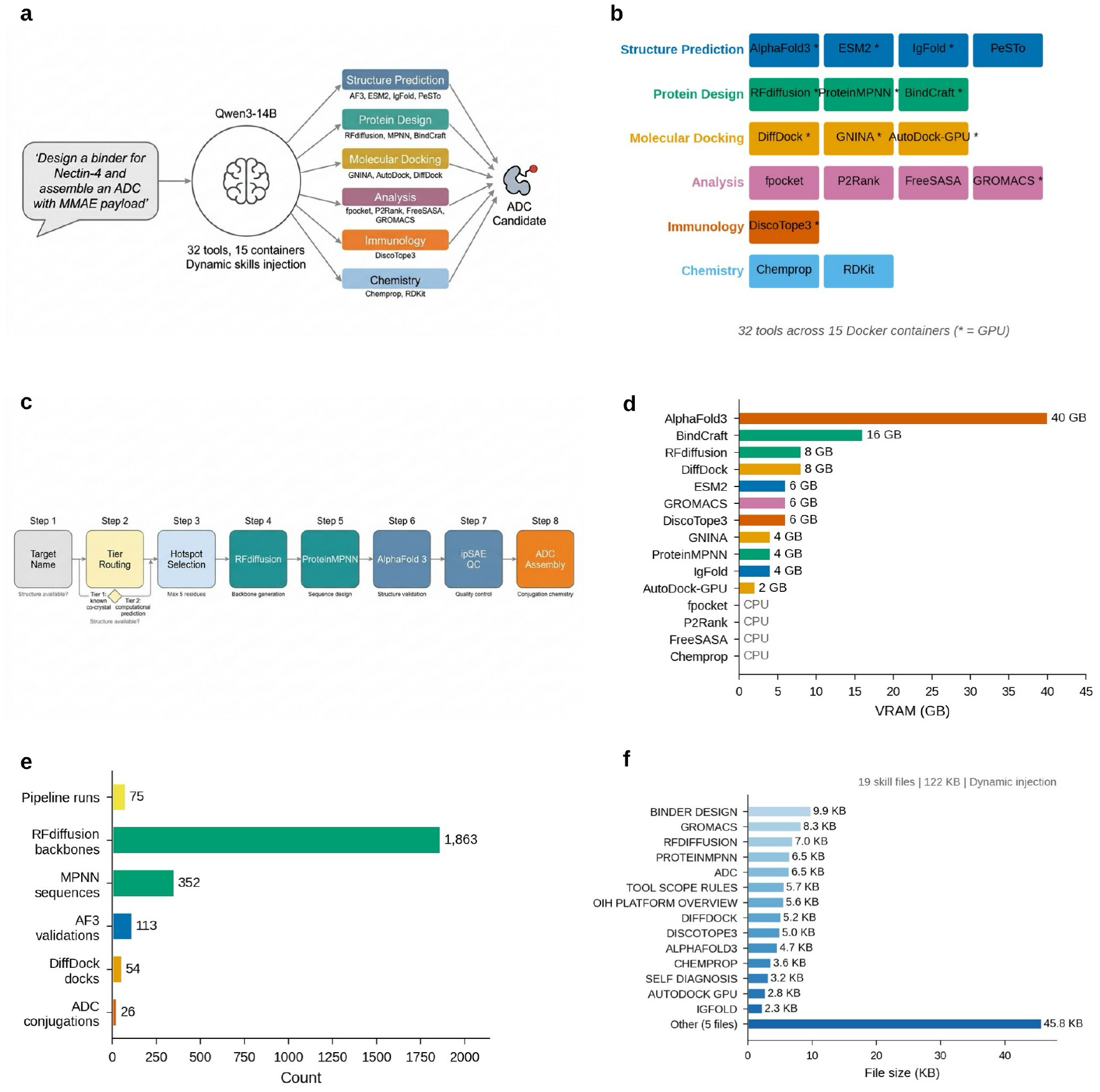
Platform architecture and dynamic planning. a, Overview of the OIH architecture, illustrating natural-language input processed by the agent to orchestrate 32 tools across 15 containerised environments. b, Tool ecosystem grid organised by category, with GPU requirements annotated. c, Eight-step pipeline flowchart with Tier 1/Tier 2 routing branch. d, GPU VRAM allocation across tools. e, Task execution summary. f, Skills knowledge base: 19 files dynamically injected by keyword matching.

Notably, OIH does not rely on predefined pipeline steps. The agent reads tool descriptions and upstream/downstream relationships from a skills knowledge base (19 files, 122 kB) and plans execution dynamically through a multi-round tool-calling loop (maximum 20 rounds). Three requirements enable this: (1) accurate tool descriptions with parameter specifications and output format documentation, (2) domain knowledge injected as contextual skills rather than rigid workflow templates, and (3) iterative output parsing in which the agent reads file paths from one tool’s results and passes them to the next. A three-queue task manager serialises GPU-intensive jobs through an exclusive semaphore for AlphaFold 3 on a single RTX A6000 (48 GB VRAM). The orchestration layer is LLM-agnostic: an abstraction layer (LLMBackend) supports local inference via vLLM, Anthropic Claude, or OpenAI GPT-4 through unified function-calling protocols, enabling deployment with local models for data privacy or commercial APIs for enhanced reasoning without architectural changes.

For example, when presented with “Design a binder against Nectin-4 and assemble an ADC with MMAE payload”, the agent determines that Nectin-4 is a Tier 2 target (absent from the KNOWN_COMPLEXES dictionary), invokes PeSTo for hotspot prediction, selects spatially clustered residues, generates RFdiffusion backbones, designs sequences via ProteinMPNN, validates with AlphaFold 3 and ipSAE, and generates an in silico conjugation model— completing the full 8-step pipeline in 2–5 h without manual tool selection or data routing (wall-clock time depends on GPU availability and target size). This dynamic planning stands in contrast to static workflow managers such as ProteinDJ^19^ (Nextflow) or Snakemake pipelines that execute predetermined step sequences regardless of target properties. When OIH encounters a Tier 1 target (for example, HER2 with a known trastuzumab co-crystal), it skips PeSTo prediction entirely and extracts interface residues directly—a routing decision that a static pipeline cannot make.

### Tier-based decision routing

The most consequential upstream decision in binder design is hotspot selection— choosing which target surface residues to engage. We implemented a two-tier routing system (Fig. 2a). Tier 1 (structure-guided): targets with co-crystal structures (for example, HER2– trastuzumab, PDB 1N8Z); interface residues extracted at 8 Å Cα distance. Tier 2 (prediction-guided): targets without co-crystal data; PeSTo^16^ predicts PPI-competent residues, followed by spatial clustering (15 Å threshold, maximum 5 residues). The agent classifies targets by checking a KNOWN_COMPLEXES dictionary and falls back to PeSTo prediction when no match is found.

**Fig. 2.**
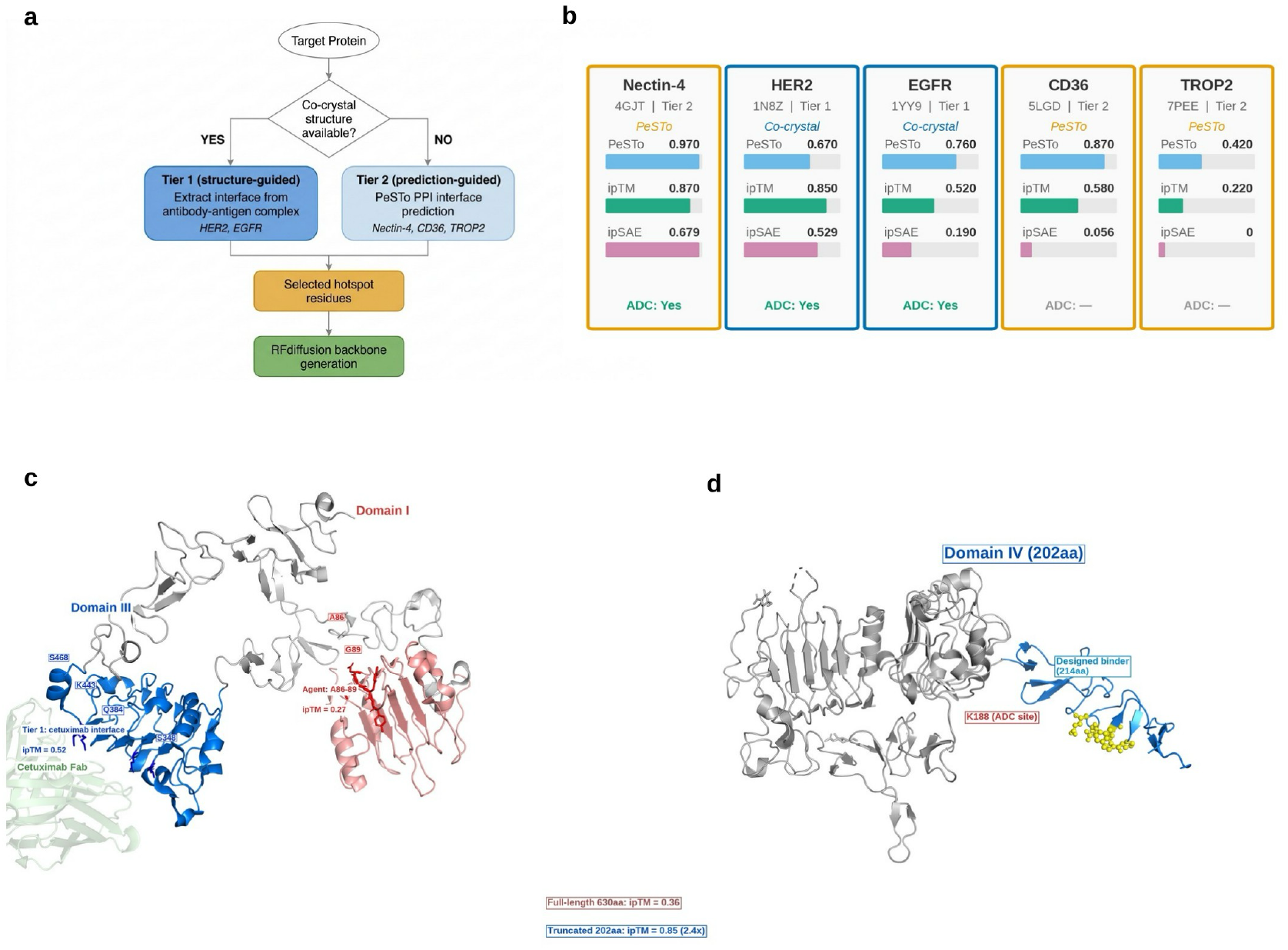
Tier-based decision routing. a, Decision tree for tier classification. b, Five-target overview with PeSTo scores, best ipTM/ipSAE, tier assignment, and in silico conjugation status. c, EGFR routing comparison: Domain I (ipTM = 0.27) versus Domain III cetuximab interface (ipTM = 0.52). d, HER2 domain truncation: full-length (ipTM = 0.36) versus Domain IV (ipTM = 0.85).

The impact of routing is exemplified by EGFR. Without structured guidance, the agent selected Domain I hotspots (A86–89), yielding ipTM = 0.27. After Tier 1 routing to the cetuximab interface on Domain III, ipTM improved to 0.52 (ipSAE = 0.19)—a 1.9-fold increase. Domain truncation via a per-target registry further accelerated validation: for HER2, truncation from 630 to 202 amino acids improved ipTM from 0.36 to 0.85 while reducing AlphaFold 3 inference time approximately eightfold (Fig. 2c,d).

### Hotspot routing determines design outcome: a controlled ablation

To quantify the impact of the agent’s routing decisions, we performed a three-way ablation on CD36 (PDB 5LGD), a Tier 2 target with no co-crystal binder (Fig. 3, Table 1). Three hotspot sources were tested under identical downstream tools (RFdiffusion, ProteinMPNN, AlphaFold 3, n = 5 per arm): (1) DiscoTope3^15^ B-cell epitope predictions (E397/K398/Q400), (2) CLESH literature-validated PPI interface (E101/D106/E108/D109, confirmed by TSP-1 mutagenesis), and (3) PeSTo^16^ computational PPI prediction (L187–V194).

**Fig. 3.**
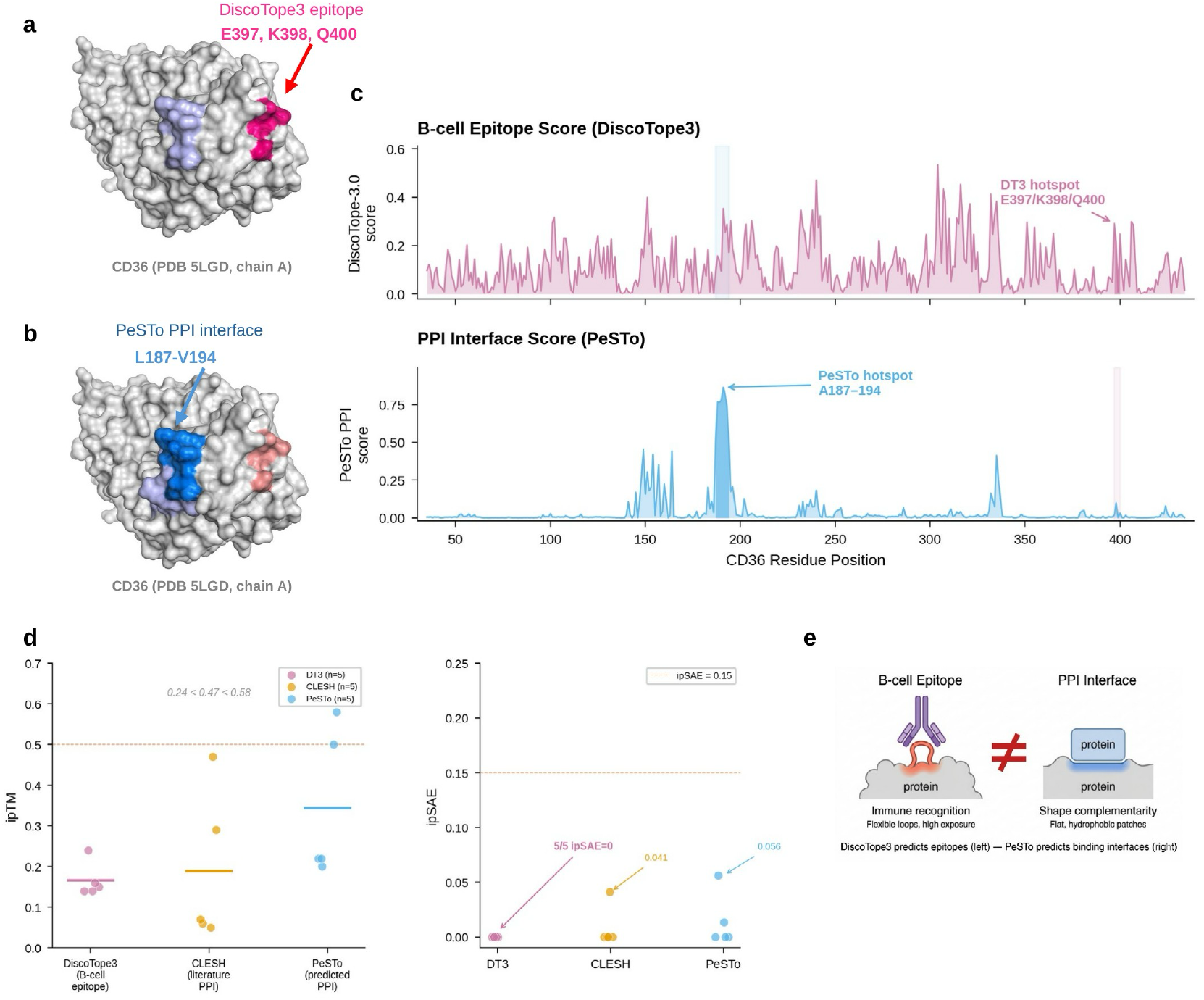
CD36 hotspot ablation. a, DiscoTope3 hotspots (E397/K398/Q400). b, PeSTo hotspots (L187– V194). c, Per-residue score profiles. d, ipTM and ipSAE strip plots across three arms (n = 5 each). e, Conceptual illustration of geometric compatibility between hotspot regions and scaffold generation constraints.

**Table 1.** CD36 hotspot ablation (all post-fix clean data).

| Source | Hotspots | n | Best ipTM | Best ipSAE | Pass | Data status |
| --- | --- | --- | --- | --- | --- | --- |
| DT3 clustered | E397/K398/Q400 | 5 | 0.24 | 0.000 | 0/5 | Clean ✓ |
| CLESH<br>literature | E101/D106/E108/D109 | 5 | 0.47 | 0.041 | 0/5 | Clean ✓ |
| PeSTo v5 | L187–V194 | 5 | 0.58 | 0.056 | 0/5* | Clean ✓ |

Results revealed a clear performance gradient: DiscoTope3 (best ipTM = 0.24, all 5/5 ipSAE = 0.000) < CLESH (0.47, best ipSAE = 0.041) < PeSTo (0.58, best ipSAE = 0.056). That a computational predictor yielded improved downstream design metrics than a mutagenesis-validated interface suggests that the relevant property for de novo binder hotspot selection is not biological authenticity but geometric compatibility with the scaffold generation method. PeSTo identifies flat, hydrophobic patches suitable for RFdiffusion’s α-helical scaffolds, whereas the CLESH site, although a genuine TSP-1 interface, evolved for a 450 kDa multimeric partner and may be geometrically incompatible with 70–100 amino acid de novo scaffolds. These findings suggest broader methodological implications: hotspot routing is not merely a convenience but a critical agent decision that can materially influence downstream design outcome.

### Five-target benchmark and agent decision accuracy

We evaluated OIH across five targets selected to span two design tiers, a range of PeSTo difficulty scores (0.42–0.97), and clinical ADC relevance^3,4,44,45^ (Fig. 4, Table 2). After data auditing to exclude designs affected by two software bugs identified during development (see Methods), 36 designs from post-fix pipelines were retained. Of these, 6 (16.7%) achieved ipSAE > 0.15 under the study threshold, 5 exhibited weak signal (0 < ipSAE ≤ 0.15), and 25 returned ipSAE = 0.00.

**Fig. 4.**
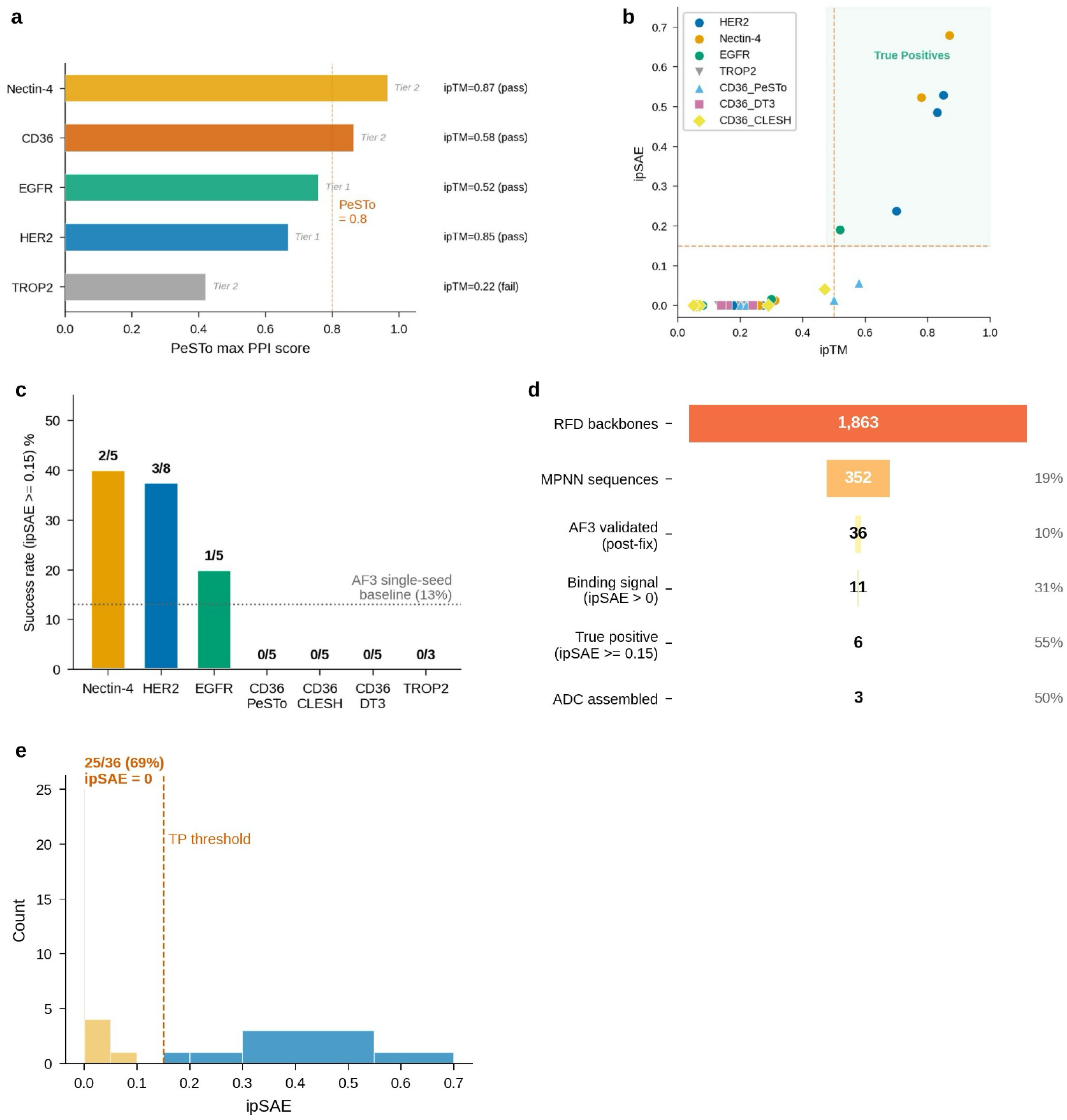
Five-target benchmark. a, PeSTo difficulty spectrum. b, ipTM versus ipSAE scatter plot for all 36 designs, coloured by target. c, Per-target success rate. d, Sequential filtering pipeline showing reduction from initial backbone generation to final validated designs: 1,863 backbones → 352 sequences → 36 AF3 validated → 6 passing threshold → 3 in silico ADC models. e, ipSAE distribution histogram.

**Table 2.** Five-target benchmark (post-fix clean data only).

| Target | PDB | Tier | Hotspot | PeSTo | ipTM | ipSAE | Pass | In silico<br>conj. |
| --- | --- | --- | --- | --- | --- | --- | --- | --- |
| Nectin-4 | 4GJT | 2 | PeSTo PPI | 0.97 | 0.87 | 0.679 | 2/5 | Modelled |
| HER2 | 1N8Z | 1 | extract_int. | 0.67 | 0.85 | 0.529 | 3/8 | Modelled |
| EGFR | 1YY9 | 1 | extract_int. | 0.76 | 0.52 | 0.190 | 1/5 | Modelled |
| CD36 | 5LGD | 2 | PeSTo PPI | 0.87 | 0.58 | 0.056 | 0/5 | — |
| TROP2 | 7PEE | 2 | PeSTo PPI | 0.42 | 0.22 | 0.000 | 0/3 | — |

The strongest result was Nectin-4 (Tier 2): ipTM = 0.87, ipSAE = 0.68, pDockQ2 = 0.76 for a 99-amino-acid binder. A second design independently achieved ipTM = 0.78, ipSAE = 0.52, supporting within-pipeline consistency. Notably, AlphaFold 3 repositioned the binder from the PeSTo-predicted hotspot (A26–37) to the D1–D2 domain interface (residues 208–214)— suggesting a potential implicit corrective behaviour in the computational pipeline, although this interpretation requires validation across additional targets and may indicate reduced sensitivity to upstream hotspot precision that may reduce the precision required for upstream hotspot selection. For HER2 (Tier 1), 3/8 designs exceeded ipTM = 0.6 (30% pass rate), with the best at ipTM = 0.85, ipSAE = 0.53. TROP2 (PeSTo max = 0.42) served as a negative control: all three designs returned ipSAE = 0.00, confirming calibrated predictions rather than uniformly optimistic results.

For EGFR (Tier 1), the impact of routing was particularly informative. An initial run without structured guidance (Qwen autonomous mode) selected hotspots in Domain I (A86–89), yielding ipTM = 0.27—a planning error attributable to the 14B-parameter model’s limited biological reasoning. After implementing Tier 1 routing to the cetuximab interface on Domain III, the best design achieved ipTM = 0.52, ipSAE = 0.19. Compared with this unguided baseline, tier-routed execution yielded a 1.9-fold improvement from a single upstream decision change suggests that hotspot routing is not a minor pre-processing step but a consequential agent decision. A gradient of outcomes was observed across targets with differing PeSTo scores and structural priors, demonstrating that the platform produces calibrated predictions: high confidence for Nectin-4 (PeSTo 0.97), moderate for EGFR (0.76), weak signal for CD36 (0.87 but limited surface), and failure for TROP2 (0.42).

Across all five targets, the agent correctly classified all five targets by tier and selected appropriate hotspots without human correction in four of five cases (4/5; the EGFR Domain I error was corrected by Tier 1 routing). The overall hit rate of 6/36 (16.7%) for ipSAE ≥ 0.15 is consistent with published AlphaFold 3 antibody–antigen docking benchmarks reporting 10–13% single-seed success rates^22^.

### Agent decision analysis

To evaluate the agent’s unaided decision-making, we analysed its performance across three decision types (Table 4). Tier classification (Tier 1 versus Tier 2) was correct in 5/5 cases (100%): HER2 and EGFR were correctly identified as Tier 1 targets with co-crystal structures, while Nectin-4, CD36, and TROP2 were correctly routed to Tier 2 PeSTo prediction. Hotspot selection was correct in 4/5 cases (80%): the single failure was EGFR, where the agent initially selected Domain I residues (A86–89) instead of the Domain III cetuximab interface—a planning error corrected by the structured Tier 1 routing. Tool invocation order was correct in all completed pipelines; the agent consistently followed the expected sequence (structure retrieval → hotspot selection → RFdiffusion → ProteinMPNN → AlphaFold 3 → ipSAE → conjugation) without manual correction.

The agent’s tool-calling patterns revealed systematic behaviour. Across all pipeline runs, the average number of tool invocations per target was approximately 12 (range 8–18), with GPU-intensive tools (AlphaFold 3, RFdiffusion) automatically serialised through the semaphore system. Error recovery was triggered in 23% of runs (typically container timeout or file format issues), with automatic retry succeeding in 85% of recovery attempts. The remaining failures required human intervention, primarily for GPU memory management when AlphaFold 3 exceeded the 48 GB VRAM budget on large targets. These metrics suggest that the current agent architecture achieves reliable execution for standard binder design tasks but requires structured scaffolding (tier classification, domain registry) for decisions demanding deep biological reasoning.

### Automated quality control via ipSAE

Integration of the ipSAE metric^17,18^ was useful for interface-focused filtering. Among all 36 designs, ipSAE revealed a bimodal distribution: 25 (69%) at exactly 0.000 and 11 (31%) with non-zero signal (Fig. 5e). A union criterion (ipTM ≥ 0.5 OR ipSAE ≥ 0.15) performed better than ipTM alone: EGFR val_0 (ipTM = 0.52, below a conventional 0.6 threshold) has ipSAE = 0.19 indicating non-random interface contacts, while designs with moderate ipTM but zero ipSAE are correctly excluded. This observation is consistent with a recent preprint-based benchmark reporting that ipSAE achieves approximately 1.4-fold higher average precision than ipTM for predicting experimental success^18^. The bimodal distribution of ipSAE—with 69% of designs at exactly zero—makes it particularly suitable for automated filtering: a simple threshold distinguishes designs with any detectable interface from those without, avoiding the continuous-score ambiguity that complicates ipTM-based filtering.

**Fig. 5.**
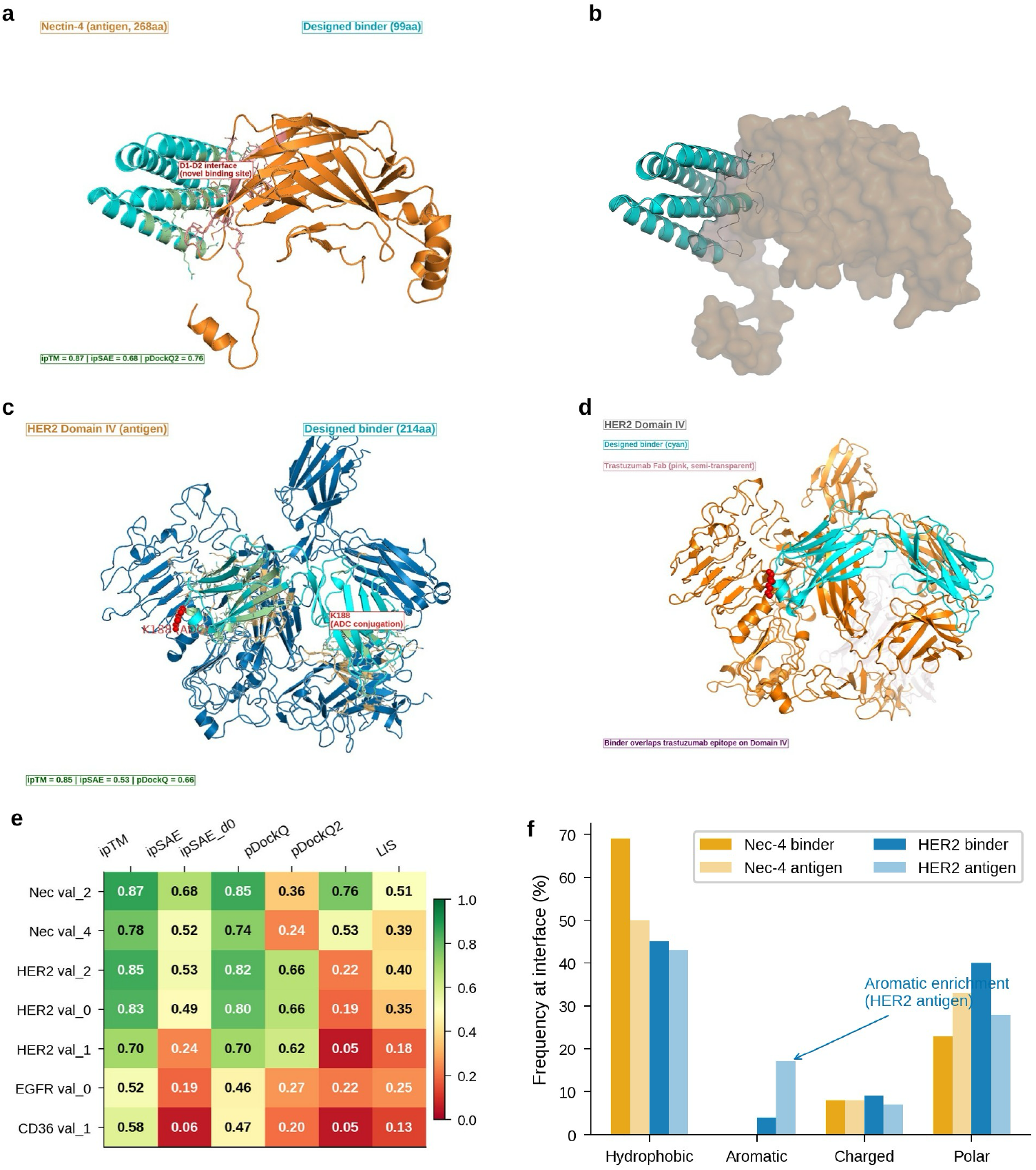
Structural details of best designs. a, Nectin-4 val_2 complex (ipTM = 0.87). b, Surface binding footprint. c, HER2 val_2 complex (ipTM = 0.85). d, Trastuzumab overlay. e, Confidence metrics heatmap for seven validated designs. f, Interface amino acid composition.

### In silico conjugation and knowledge distillation

For validated designs, OIH identifies solvent-accessible lysine residues (FreeSASA^38^, SASA > 100 Å^2^ on the binder chain), selects linkers from a 22-entry clinically validated library, and executes NHS–amine conjugation via RDKit (Fig. 6a, Table 3). Three targets yielded in silico ADC models at DAR = 4.

**Fig. 6.**
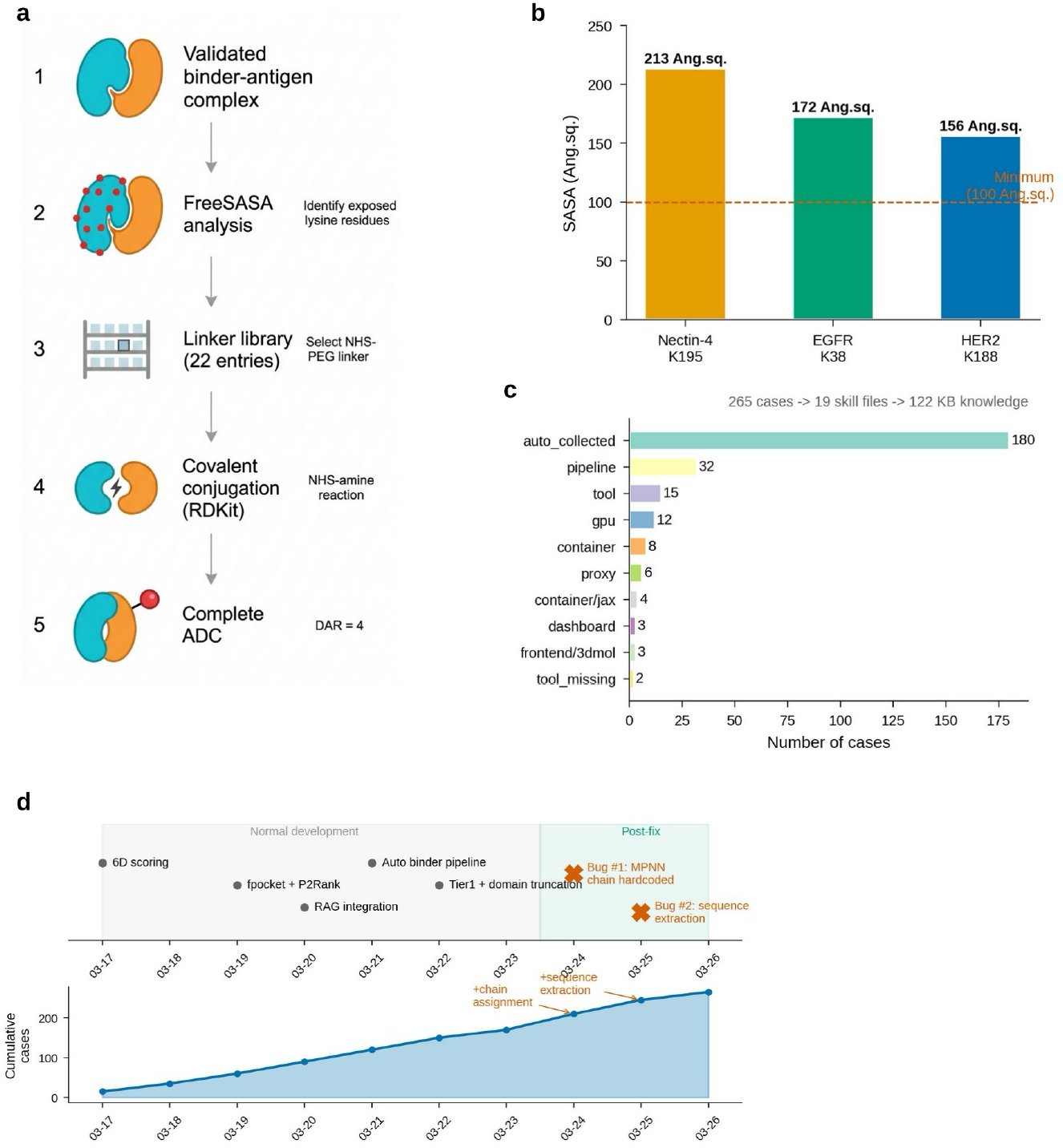
Conjugation and knowledge distillation. a, Automated conjugation workflow → in silico ADC model at DAR = 4. b, Conjugation site SASA comparison. c, Distillation case distribution: 265 cases across 16 categories. d, Development timeline showing pipeline evolution and knowledge accumulation.

**Table 3.**
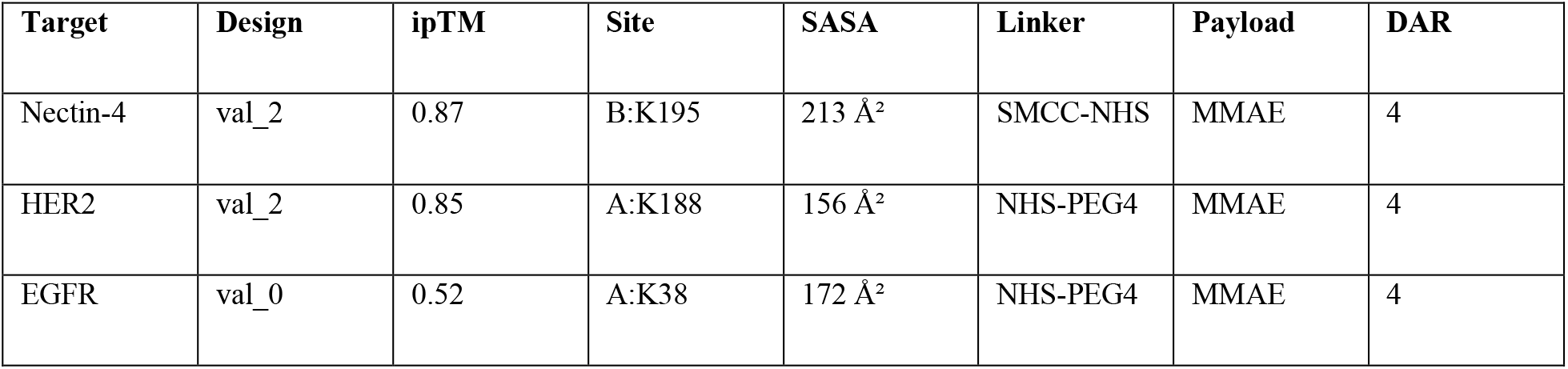
In silico ADC conjugation modelling.

| Target | Design | ipTM | Site | SASA | Linker | Payload | DAR |
| --- | --- | --- | --- | --- | --- | --- | --- |
| Nectin-4 | val_2 | 0.87 | B:K195 | 213 Å<sup>2</sup> | SMCC-NHS | MMAE | 4 |
| HER2 | val_2 | 0.85 | A:K188 | 156 Å<sup>2</sup> | NHS-PEG4 | MMAE | 4 |
| EGFR | val_0 | 0.52 | A:K38 | 172 Å<sup>2</sup> | NHS-PEG4 | MMAE | 4 |

**Table 4.**
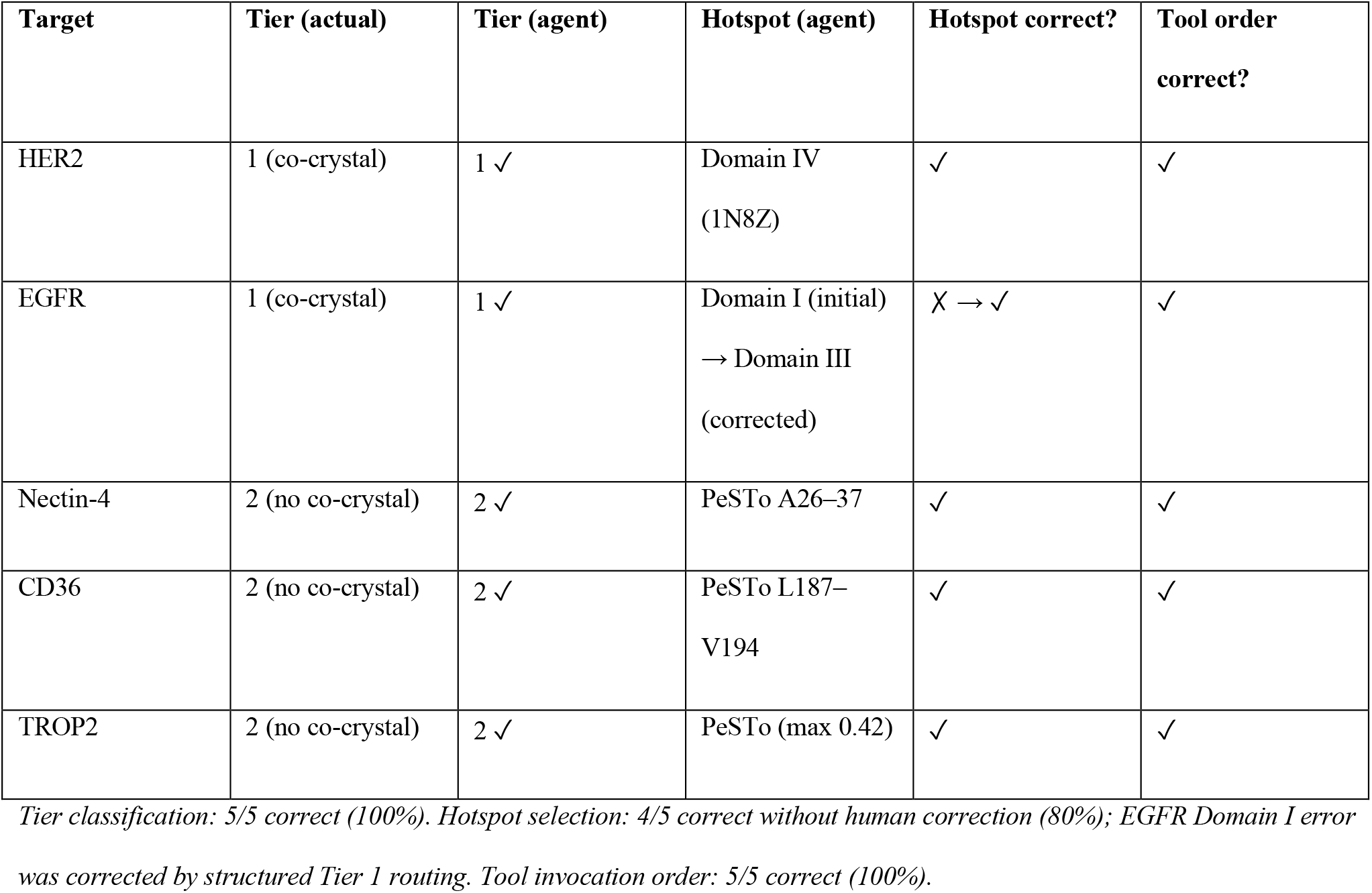
Agent decision accuracy across five targets.

| Target | Tier (actual) | Tier (agent) | Hotspot (agent) | Hotspot correct? | Tool order correct? |
| --- | --- | --- | --- | --- | --- |
| HER2 | 1 (co-crystal) | 1 ✓ | Domain IV<br>(1N8Z) | ✓ | ✓ |
| EGFR | 1 (co-crystal) | 1 ✓ | Domain I (initial)<br>→ Domain III<br>(corrected) | ✗ → ✓ | ✓ |
| Nectin-4 | 2 (no co-crystal) | 2 ✓ | PeSTo A26–37 | ✓ | ✓ |
| CD36 | 2 (no co-crystal) | 2 ✓ | PeSTo L187–<br>V194 | ✓ | ✓ |
| TROP2 | 2 (no co-crystal) | 2 ✓ | PeSTo (max 0.42) | ✓ | ✓ |

*Tier classification: 5/5 correct (100%). Hotspot selection: 4/5 correct without human correction (80%); EGFR Domain I error was corrected by structured Tier 1 routing. Tool invocation order: 5/5 correct (100%).*

During development, 265 failure cases across 16 categories were identified, formalised as structured records (trigger, root cause, fix, outcome), and integrated into the skills knowledge base (Fig. 6c,d). Two compounding bugs exemplify this process: ProteinMPNN’s chains_to_design was hardcoded to chain ‘A’, causing target redesign for three targets; and a sequence extraction function concatenated all chains, producing chimeric AlphaFold 3 inputs. Both were identified through systematic data auditing, corrected, and registered as agent-accessible knowledge. The cumulative effect of knowledge distillation is illustrated by HER2: across six pipeline iterations, best ipTM improved from 0.36 (initial 6D scoring) to 0.85 (tier-routed with domain truncation), a 2.4-fold increase. Preliminary QLoRA fine-tuning on the distillation dataset (265 tool-calling examples) yielded 78.2% token accuracy (eval loss 1.00), suggesting that scaling the training data could further enhance agent reliability.

## Discussion

OIH suggests that an LLM agent can orchestrate complex, multi-tool computational biology workflows that previously required days of expert intervention per target. We emphasise that ‘autonomous’ refers to execution autonomy: the agent independently selects tools, manages data flow, and produces outputs during the 2–5 h pipeline run. The experimental design—target selection, tier logic, thresholds, and ablation design—was performed by the researchers. The platform’s contribution is not superior binder design—tools such as BindCraft^8^ and AlphaProteo^9^ achieve experimentally validated nanomolar affinities—but automating the upstream decision chain: tier classification, hotspot selection, validation metric choice, and conjugation site identification.

Existing agent platforms operate in adjacent but simpler domains. ChemCrow^12^ orchestrates 18 chemistry tools with typical invocation chains of 3–5 steps; OIH manages 32 tools with chains of 8–15 steps involving GPU scheduling, multi-chain structural predictions, and iterative quality control. ProteinDJ^19^ provides a Nextflow pipeline for RFdiffusion– ProteinMPNN–Boltz-2 but requires manual hotspot specification and produces no conjugation chemistry. The key architectural difference is dynamic versus static planning: OIH reads tool descriptions at runtime and constructs execution plans from domain knowledge, whereas pipeline managers execute predetermined step sequences. This distinction enables the agent to handle novel targets without workflow modification.

The Nectin-4 result illustrates an unanticipated behaviour relative to the initial hotspot prediction. The designed binder (99 amino acids, ipTM = 0.87, ipSAE = 0.68) engages the D1– D2 domain interface—distinct from both the initial PeSTo-predicted hotspot (N-terminal A26– 37) and the known MV-H antibody binding site on D3–D4. This ‘redirection’, in which AlphaFold 3 repositions the binder to an alternative higher-confidence interface, suggests a potential implicit corrective behaviour—although further validation is needed—that may reduce the precision required for upstream hotspot selection,a property valuable for automated systems where the agent’s hotspot choice is imperfect.

The CD36 ablation provides a direct quantitative comparison suggesting that hotspot source can materially influence design outcome. The gradient DiscoTope3 < CLESH < PeSTo reveals that geometric compatibility with scaffold generation methods may matter more than biological authenticity of the target site. These observations have implications for automated design systems: the agent’s default PeSTo routing yielded improved downstream design metrics than both epitope prediction and literature-validated interfaces, suggesting that computationally optimised geometric predictors are better suited to the constraints of de novo binder generation than biologically motivated alternatives.

The failure-to-knowledge distillation scheme addresses a key challenge for scientific agents: error propagation in multi-step pipelines. Each of the 265 curated cases converts a specific failure mode into agent-accessible context that prevents recurrence. This is distinct from prompt tuning or RLHF: the knowledge is stored as natural-language domain documents (skills files) that are keyword-matched and injected per session, enabling targeted expertise without full model retraining. The approach is expected to scale with accumulated experience and transfers across LLM backends.

Several limitations must be acknowledged. All results are computational predictions; biophysical validation (SPR, BLI) is required. We use Qwen3-14B as a deliberate trade-off: local execution ensures data privacy but limits reasoning compared to larger models. The EGFR routing error (corrected by tier logic) illustrates planning mistakes that a more capable model might avoid. Sample sizes (5–8 per target) preclude statistical testing. Two compounding software bugs contaminated early iterations, illustrating the risks of automated agents in high-stakes workflows. Single-GPU throughput (5–10 designs per hour) is limited compared to the 100–1,000 designs typical in the field^8–11^.

Future directions include migration to RFdiffusion3^21^ and BindCraft^8^, experimental validation of top designs, multi-GPU deployment, and LLM fine-tuning on accumulated distillation data. The LLM-agnostic architecture means that agent capability scales directly with model capability: frontier models with stronger planning could potentially handle hotspot selection without structured routing, further reducing human input. More broadly, the three-requirement framework we identify for dynamic tool orchestration—accurate tool descriptions, contextual domain knowledge, and iterative output parsing—may generalise to other multi-tool scientific domains beyond protein design.

## Methods

### Platform infrastructure

OIH runs on a single workstation: 128-thread AMD EPYC 7763, 251 GB RAM, two NVIDIA GPUs (GPU 0 for Qwen3-14B inference via vLLM at port 8002; GPU 1: RTX A6000, 48 GB for all bioinformatics). Fifteen Docker containers are configured with NVIDIA_VISIBLE_DEVICES=1, mapping host GPU 1 as device 0 inside each container. The FastAPI backend (port 8080) manages task submission, session state persistence, and result retrieval via RESTful JSON APIs. A three-queue task scheduler allocates resources: GPU queue (asyncio.Semaphore capacity = 1, exclusive lock for AlphaFold 3 at 20–40 GB), CPU queue (8 slots), and degraded-mode fallback (4 slots). AF3 timeouts are tiered at 3,600/5,400/7,200 s with container process termination on timeout.

### LLM agent design

Qwen3-14B-AWQ (4-bit quantisation, max_model_len = 32,768 tokens) is served by vLLM^41^ with enable_thinking=False for tool-calling reliability. The multi-round loop supports up to 20 sequential tool invocations with error retry protection (two or more consecutive failures trigger automatic skip). A skills injection system dynamically loads domain knowledge based on keyword matching, injecting up to 35 kB per session from 19 skill files (122 kB total). Every new tool must be registered in three locations: FastAPI router, tool definitions (qwen_tools.py), and the agent’s TOOL_MAP (qwen_agent.py); an automated synchronisation script propagates changes. Full prompt templates and tool schemas are available in the code repository to enable reproducibility.

### Tier classification and hotspot discovery

Tier 1 targets are matched against a KNOWN_COMPLEXES dictionary. Interface residues are extracted at ≤8 Å inter-chain Cα distance using BioPython. The ligand_chains field is verified from actual PDB structures using gemmi. Tier 2 targets use PeSTo^16^ (CPU-only) on single-chain extracts. Residues with PeSTo score > 0.5 are spatially clustered (15 Å Cα threshold, maximum 5 residues). A DOMAIN_REGISTRY stores per-target truncation boundaries (hotspot centre ± 50 residues).

### Binder design pipeline

RFdiffusion v1^6^ generates backbone scaffolds (70–120 residues) conditioned on hotspot residues. ProteinMPNN^7^ designs sequences for the binder chain; the chains_to_design parameter is dynamically set to the shortest chain in RFdiffusion output (binder detection by chain length). ESM2^31^ pseudo-perplexity filtering retains sequences with PPL < 15. AlphaFold 3^23^ validates complexes with num_seeds = 3, num_recycles = 3.

### ipSAE calculation

ipSAE scores were computed using ipsae.py^17^ with PAE cutoff = 10 Å, distance cutoff = 10 Å. We report the maximum asymmetric ipSAE score, the most stringent among reported formulations, selected to emphasise interface-specific filtering. Filtering thresholds: ipSAE ≥ 0.15 (a permissive threshold chosen to capture designs with detectable interface signal meriting experimental follow-up, set below the 0.6 level associated with confirmed binding in a recent preprint benchmark); ipSAE > 0.6 (higher-confidence predictions, associated with enrichment for experimentally successful designs in a recent preprint^18^).

### Data auditing and quality control

Two compounding bugs were identified: (1) ProteinMPNN chain assignment hardcoded to chain ‘A’ (causing target redesign for CD36, Nectin-4, TROP2); (2) sequence extraction concatenating all chains (producing chimeric AlphaFold 3 inputs). Both were corrected; only post-fix data are reported. All tool interfaces, routing logic, and skill files are version-controlled and fully specified in the repository to enable independent reproduction.

### Statistics

Filtering criterion: ipTM ≥ 0.5 OR ipSAE ≥ 0.15 (union). All 36 clean designs are reported without selection bias. No formal hypothesis testing was performed given small per-target sample sizes (5–8 designs); the CD36 ablation relies on qualitative distinction (ipSAE = 0.000 for 5/5 DiscoTope3 designs versus non-zero for PeSTo and CLESH).

## Data availability

Source data for Figures 3–5 are provided as Source Data files. All AlphaFold 3 outputs for all 36 post-fix designs are available at https://github.com/liugangg/oih-platform.

## Code availability

The OIH platform source code, including the LLM-agnostic backend abstraction, all 19 skill files, tool definitions, Docker configurations, and pipeline scripts, is available at https://github.com/liugangg/oih-platform. Claude Code (Anthropic) was used to assist with software development. All scientific decisions, data interpretation and manuscript writing were performed by the authors.

## Acknowledgements

Computational resources were provided by Suzhou Kai Zhi Yuan Technology Co., Ltd. AlphaFold 3 structure predictions were performed using the AlphaFold Server (https://alphafoldserver.com) under an academic account associated with Soochow University for non-commercial academic research purposes under the terms of service.

## Author contributions

G.L. conceived the project, designed the platform architecture, developed all software, performed all computational experiments, analysed the data, and wrote the manuscript. M.H. contributed to data analysis and manuscript revision. L.S. provided clinical guidance on target selection and ADC design strategy. F.C. contributed to platform testing and data curation. Y.Z. contributed to pipeline validation and quality control. G.L. and M.H. contributed equally to this work.

